# Bumblebees develop more efficient traplines than honey bees

**DOI:** 10.1101/2020.12.22.423907

**Authors:** Alexis Buatois, Thibault Dubois, Mathieu Lihoreau

## Abstract

Central place foraging pollinators, such as bees, tend to learn multi-destination routes (traplines) to efficiently visit known feeding locations and return to their nest. To what extent these routing behaviours are shared across species is unknown. Here we ran laboratory experiments to compare trapline formation and efficiency by foragers of two social bee species that differ in their collective foraging strategies: the solo foraging bumblebee *Bombus terrestris* and the mass foraging honey bee *Apis mellifera*. In a simple routing task with four artificial flowers, both bumblebees and honey bees developed a stable route, although honey bees were slower and less efficient to do so. In a more complex routing task with six flowers, only bumblebees developed a stable route. Honey bees took a longer time to discover all flowers and never integrated them in a single route. Simulations of a model of trapline formation show that these inter-specific differences can be replicated by adjusting the strength of a single learning parameter. Comparing bumblebees and honey bees in the same experimental conditions thus revealed key differences in their spatial foraging strategies, potentially driven by social constraints.

## Introduction

Central place foraging pollinators often develop multi-destination routes (or traplines) to efficiently exploit patchily distributed feeding sites (Thomson et al. 1996). Trapline foraging is widespread in nectar feeding insects (orchid bees: Janzen 1971; bumblebees: Thomson et al. 1997; honey bees: Buatois and Lihoreau 2016; butterflies: Gilbert and Singer 1975) but also in birds (hummingbirds: Tello-Ramos et al. 2015) and mammals (bats: Lemke 1984; opposums: Wooller et al 1999). This routing behaviour is thought to optimize the exploitation of plants resources, by adjusting the timing of nectar collection and deterring competitors (Possingham 1989; Ohashi and Thomson 2005).

Trapline formation has been best studied in bumblebees using arrays of artificial flowers (i.e. feeders) in the lab and in the field (Ohashi and Thomson 2009; Lihoreau et al. 2013). Individual bumblebees given an access to a small number of flowers for several hours often find and use the shortest sequence to visit all flowers once and return to the nest (Ohashi et al. 2007; Lihoreau et al. 2010, 2012a). Analyses of the flight paths of bumblebees between visiting flowers, show that insects also tend to optimize travel distances (Lihoreau et al. 2012b; Woodgate et al. 2017). Trapline formation can be modelled using an iterative improvement heuristic of vector navigation replicating trial-and-error learning by bees (Lihoreau et al. 2012b; Reynolds et al. 2013). In this approach, the bumblebee compares the net length of all the route segments (i.e. straight movements between two flowers or between a flower and the colony nest) composing the route it has just experienced, and increases its probability to reuse these segments in future if the new route is shorter (or of the same length) than the shortest route experienced before.

Recently, honey bees were also reported to be capable of developing traplines between four artificial flowers (Buatois and Lihoreau 2016). The indirect comparison of the foraging patterns of bumblebees and honey bees, using network statistics on data obtained in different studies, suggests that honey bees develop less efficient routes and are less faithful to these routes than bumblebees (Pasquaretta et al. 2017). This may be explained by major differences in the social ecology of these bees. Bumblebees, as solo foragers, primarily rely on the acquisition of individual information to locate and exploit plant resources (Dornhaus and Chittka 1999). By contrast, honey bee are mass foragers and use social information to collectively exploit profitable food resources (e.g. they use the waggle dance for recruiting nestmates to resources 100 m away from the nest, von Frisch 1967). Honey bees may therefore invest less in individual sampling and spatial learning than bumblebees (Buatois and Lihoreau 2016). Further understanding these behavioural differences could have considerable implications for assessing the impact of foraging strategies and social structure on the evolution of cognition in two major pollinators and model species in for insect behaviour (Farris 2016; Traniello et al. 2019). Such approach requires to study the animals in strictly identical setups and make quantitative comparisons of their performances (Chittka et al. 2012).

Here, we tested the hypothesis that honey bees have poorer abilities to develop foraging routes than bumblebees, by comparing the spatial patterns of individuals of both species in the same arrays of artificial flowers in an indoor flight room. Since the number of possible routes to visit each flower once and return to the nest increases factorially with the numbers of flowers to visit (Lihoreau et al. 2013), we tested the influence of task complexity by observing bumblebees and honey bees foraging in arrays of four flowers (24 possible routes) and six flowers (720 possible routes). We then compared the performances of foragers of both species with numerical simulations of a model of trapline formation.

## Materials and methods

### Bees

We used a small colony (a queen and ca. 2000 workers) of honey bees (*Apis mellifera*, Buckfast) originating from the experimental apiary of the University Paul Sabatier – Toulouse III (France). We used a normal size colony (a queen and ca. 200 workers) of bumblebees (*Bombus terrestris*) from Biobest (Belgium). Both honey bee and bumblebee hives were equipped with a transparent entrance tube fitted with gates to precisely control the traffic of foragers. Colonies were provided *ad libitum* defrosted pollen directly into the hives. Workers collected sucrose solution (40% w/w in water) from artificial flowers in the flight room.

### Flight room

We conducted the experiments in an indoor flight room (length: 7 m, width: 5 m, height: 3 m; **Fig.S1A**) equipped with 12 wide spectrum LED lights (6500K, Phillips, The Netherlands) replicating natural sunlight and daily photocycle (15h Light/ 9h Dark). Four posters uniquely characterized by a bicolored pattern were placed on the room walls to provide 2D visual landmarks to the bees (**Fig.S2**).

### Artificial flowers

Each flower consisted of a 6 cm diameter blue landing platform sitting on a 10 cm high transparent plastic cylinder. The cylinder was attached to a clamp stand 50 cm above ground (**Fig.S1B**, Buatois and Lihoreau 2016). A yellow mark in the middle of the landing platform indicated the location of a controlled volume of sucrose solution (40% w/w, range: min 20 µl – max 140 µl) dispensed using an electronic micropipette (Brand Handystep 705000, Germany).

### Flower arrays

Bees were tested either in a four-flowers array or in a six-flowers array (see flower coordinates in **Fig.S1A**). We used these two arrays to provide bees with spatial tasks of increasing complexity. In the four-flowers arrays there was 24 possible sequences to visit all flowers once starting and ending at the colony nest entrance (including the two optimal sequences F1-F2-F3-F4 or F4-F3-F2-F1; Figs 1 and S1A). In the six-flowers array, there were 720 possible sequences (including the two optimal sequences: F1-F2-F3-F4-F5-F6 or F6-F5-F4-F3-F2-F1; Figs 3 and S1B). Flower arrays were generated using a computer program (R code available in Text S1) designed to maximize the discrepancy between the two optimal flower visitation sequences minimizing travel distance to visit all flowers (e.g. four-flowers array: F1-F2-F3-F4 or F4-F3-F2-F1) and the flower visitation sequence linking all unvisited nearest neighbour flowers (e.g. four-flowers array: F1-F4-F5-F2) given the dimensions of our flight room. The distance between neighbour flowers ranged from 1.48 m to 4.19 m in the four-flowers array, and 1.13 m to 4.22 m in the six-flowers array. Since bee workers detect visual targets from a background subtending a minimum visual angle between ca. 3° (*B. terrestris;* Dyer et al. 2008*) and* 5° (*A. mellifera;* Giurfa et al. 1996; Kapustjansky et al. 2010), we assumed that bees could visually detect the 50 cm tall flowers from any location in the flight room. The small spatial scale of the experimental arrays (i.e. the maximum length between a flower and the nest was 6.41m) prevented any dance communication between the honey bees (von Frisch 1967).

**Figure 1:**
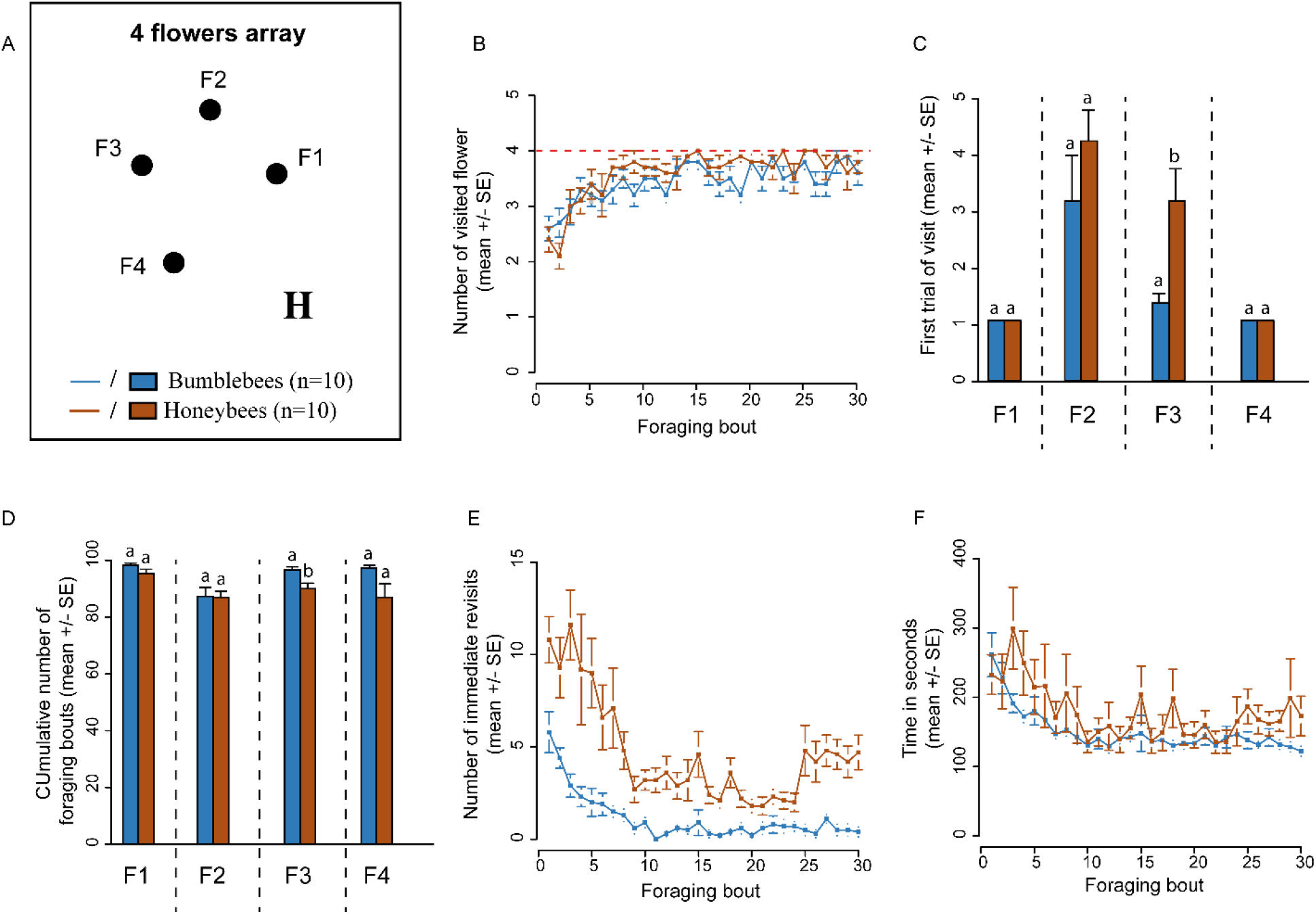
Route formation in the four-flowers array. **(A)** Schematic representation of the flower array. Flowers are labelled F1-F4. H is the colony hive location. **(B)** Number of flowers visited per foraging bout. **(C)** Cumulated number of foraging bouts before the flower was first visited. **(D)** Percentage of foraging bouts for which each flower was visited. **(E) N**umber of immediate revisits to flowers per foraging bout. **(F)** Flight duration per foraging bout. The mean ± standard error is presented for each graph. Different letters indicate a significant difference in **(C)** and **(D)** (Wilcoxon Rank-test for independent data).

### Traplining experiments

We pre-trained the bees collectively on an artificial flower delivering *ad libitum* sucrose solution (3.5cm petri dish full of sucrose solution on top of blue landing platform, 40% w/w) and marked them with acrylic paint on the thorax for individual identification (von Frisch 1967). Once a bee made regular foraging bouts (foraging trips starting and ending at the colony hive entrance), we measured its nectar crop capacity (average ± SE; bumblebees: 120±20 µL, N = 20; honey bees: 42.3±10 µL, N = 19) by estimating the average total amount of sucrose solution collected by that bee over three foraging bouts (Lihoreau et al. 2010). We then tested the bee either for 30 consecutive foraging bouts in the four-flowers array or 50 consecutive foraging bouts in the six-flowers array. The number of trials differed between the two arrays based on previous observations that bees require more time to develop routes between six flowers than in a more simple array of four flowers (Lihoreau et al. 2010, 2012a). Each flower provided either 1/4th (four-flowers array) or 1/6th (six-flowers array) of the bee’s nectar crop capacity and was refilled with sucrose solution between each foraging bout. Between testing different bees, flowers were cleaned with ethanol (70% w/w) to remove chemical cues that could influence the next foragers (Giurfa and Núñez 1992; Pearce et al. 2017). In total, 10 bumblebees and 10 honey bees were tested in the four-flowers array, and another 10 bumblebees and 9 honey bees were tested in the six-flowers array (N = 39 bees in total). All data were collected by an experimenter using the software Ethom (Taiwanica 2000). For each foraging bout of each bee, we recorded the time when the bee left the hive, each time it landed on a flower, took off, and the time when it returned to the nest.

### Modeling

We compared the observational data to numerical simulations of an agent-based model of trapline development (see details in Lihoreau et al. 2012b; Reynolds et al., 2013). Briefly, in the model the agent (bee) forages according to the following set of rules: (1) the bee has a probability of using movement vectors joining two targets (two flowers, or the nest and a flower); (2) the initial probability of using a vector depends on the distance between the two targets (probabilities are inversely proportional to the squared distance between targets and are normalized with respect to all targets); (3) the bee computes the net length of the route travelled by summing the lengths of all vectors comprising the route; (4) if the route passed through all the flowers at least once (thus filling the nectar crop capacity of the bee), the bee compares the net length of the current route to the net length of the shortest route experienced so far that passes through all the flowers; (5) if the length of the new route is inferior or equal, the probabilities of using the vectors comprising this new route in the next foraging bout are multiplied by a common factor (learning factor, lf) and all probabilities are rescaled with respect to all flowers so that they sum to unity. (6) If the bee returns to the nest, the foraging bout ends no matter the number of flowers visited. Revisits to flowers are not taken into account in the learning process, so that vectors including revisits to any flower are not reinforced through learning. We explored outputs of the model with values of learning factors ranging between 1 and 2, as suggested by Reynolds et al. (2013) for simulations at small spatial scales. A learning factor of 1.0 replicates situations where the bee does not learn (i.e. null model). Increasing values of the learning factor should lead bees to converge faster towards a stable optimized trapline. For each value of learning factor, we simulated 1000 times the execution of 30 foraging bouts (for the four-flowers array) or 50 foraging bouts (for the six-flowers array).

### Data analyses

We ran all analyses in R (R Development Core Team 2016).

#### Behavioural data

From the behavioural data, we extracted raw sequences of flower visitation (see Table S1-4). We calculated the total duration of a foraging bout (time between departure and arrival at the hive), the number of different flowers visited per foraging bout, the number of foraging bouts for which each flower has been visited during the whole experiment, and the number of immediate revisits to the same flowers per foraging bout (no other flower visited in between). A bee was considered to use a route (i.e., flower visitation sequence without immediate and non-immediate revisits) significantly more often than expected by chance if the route was used at least four times in the four-flowers array (24 possible routes in total), and two times in the six-flowers array (720 possible routes in total). The ‘primary route’ for one bee was its most used one according to these previous rules (Lihoreau et al. 2010).

We assessed route optimality using an index of route quality (i.e., the squared number of different flowers visited divided by the length of the route). The index of each route was divided by the route quality of the optimal route, so that its value varied between 0 (very poor route quality) and 1 (quality equal to the optimal route). We assessed route repeatability using a determinism index (DET; see details in Ayers et al., 2015). This index is set between 0 (no repeated visitation patterns in a visitation sequence) and 1 (complete traplining). We computed the DET for groups of 10 successive foraging bouts, using a minimum length of recurrence of 4 and without considering perpendicular diagonals.

We compared the behavioural performance (number of flowers visited, number of immediate revisits and time flying per foraging bout) of bumblebees and honey bees using generalized linear mixed models (GLMMs) with foraging bouts within bee identity as a random factor using lme4 package (Bates et al., 2014). Regarding the first trial each flower was visited, as well as the percentage of bout each flower was visited, the comparison between species within each individual flower was done using a Wilcoxon Rank-test for independent data.

#### Model simulations

For each model simulation (set of 1000 stimulations), we extracted the visitation sequences, the number of different flowers visited per foraging bout, the travel distance for each foraging bout, the route quality and the DET. We compared the behavioural data to the model simulations using GLMMs to check for significant differences in route quality and DET. We used the statistical differences in slopes (seen in the model as the interaction between the bout and treatment effects) to discriminate the experimental data and the models. A similar statistical comparison for treatment effect alone can be found in the supplementary materials (Table S5).

## Results

### Four-flowers array

We first compared the flower visitation sequences of bumblebees and honey bees in an array of four flowers (**Fig.1A**). Bumblebees and honey bees similarly increased the number of different flowers visited per foraging bout as they gained experience with the array (**Fig.1B;** Poisson GLMM; species, est=0.02, z_595_=0.19, p=0.85; foraging bout: est=0.007, z_595_=2.12, p=0.03; species x foraging bout: est=0.002, z_595_=0.32, p=0.75). Foragers of both species discovered the four flowers during their first five foraging bouts, but honey bees were significantly slower to find the third flower that was the closest to the door in the experimental room (**Fig.1C;** Wilcoxon test; *F3*: U=19.5, p=0.01, p>0.05 for all other flowers). Bumblebees and honey bees visited all flowers in more than 92.4% ± 2.5 of their foraging bouts, although honey bees visited significantly less the third flower (**Fig.1D;** Wilcoxon test; *F3*: U=83.5, p=0.01, p>0.05 for all other flowers).

Bumblebees and honey bees similarly decreased their number of immediate revisits to flowers per foraging bout through time (**Fig.1E;** Poisson GLMM; foraging bout: est=-0.1, z_595_=-12.8, p<0.0001). Foragers of both species also gradually decreased their flight duration (**Fig.1F**; Poisson GLMM; foraging bout: est=-0.01 z_595_=-29.35, p<0.0001). However, honey bees showed a lower reduction of immediate revisits (**Fig.1E**; species x foraging bout: est=0.05, z_595_=6.43, p<0.0001) and spent consequently more time flying in search for the four flowers (**Fig.1F**; species x foraging bout: est=0.002, z_595_=3.81, p=0.0001) than bumblebees. As a result, honeybees showed a significantly lower increase of route quality with time (Binomial GLMM; foraging bout: est=0.163, z_595_=5.720, p<0.0001, species x foraging bout: est=-0.137, z_595_=-4.353, p<0.0001; Fig. 5A).

All bumblebees and honey bees used a primary route (i.e. most often used flower visitation sequences excluding revisits) by the end of the experiment (**Fig.2A, B**). The optimal sequence (F1-F2-F3-F4 or F4-F3-F2-F1) was used as a primary route by 30% of the bumblebees (3 out of 10) and 80% of the honeybees (7 out of 10). On average, bumblebees used their primary route in 36.7 ± 4% (SE) of their foraging bouts, and honeybees in 33 ± 5% (SE) (**Fig.2**). Foragers of both species first used this route within their 10 first foraging bouts (bumblebees: 7± 1 bout, honeybees: 8 ± 1 bout; Wilcoxon-rank test; U=42, p=0.56) and showed a similar increase in route similarity (DET) with time (Binomial GLMM; foraging bout: est=0.223, z_415_=2.493, p=0.013; honey bee x foraging bout: est=0.301, z_415_=0.782, p=0.4341; Fig. 5A).

**Figure 2:**
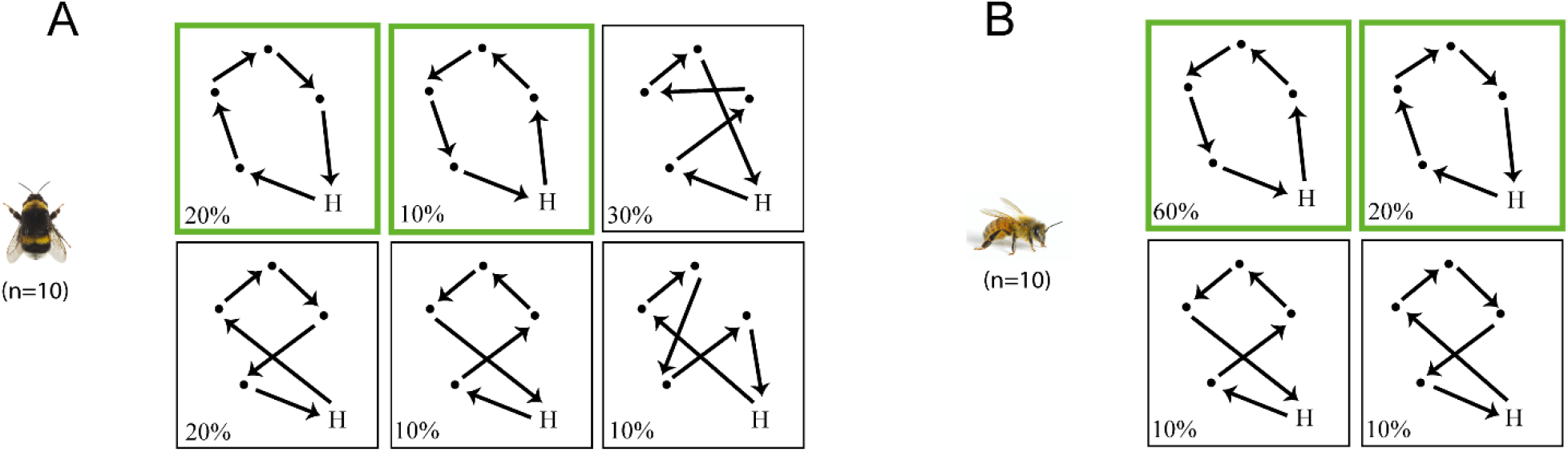
Primary routes in the four-flowers array. **(A)** Primary route used by bumblebees. **(B)** Primary route used by honeybees **(B)**. Black dots represent flower locations. H is the colony hive location. Arrows indicate the direction in which the bee moved. The percentage indicates the proportion of bees that used this route pattern. Optimal routes are highlighted in green.

Thus, in the four-flowers array, all bumblebees and honey bees developed a route between the four flowers. Honey bees were slower to locate all flowers and to reduce their revisits to empty flowers, even though foragers of both species converged towards repeatable efficient routes towards the end of the experiment.

### Six-flowers array

To test whether the complexity of the spatial problem had an influence on the routing performances of bees, we compared the flower visitation sequences of bumblebees and honey bees in an array of six flowers (**Fig.3A**).Bumblebees and honey bees increased the number of different flowers visited per foraging bout as they gained experience with the array (**Fig.3B**; Poisson GLMM; foraging bout: est=0.003, z_945_=2.16, p=0.03; species x foraging bout: est=-0.001, z_945_=-0.52, p=0.6). However, honey bees visited significantly less flowers per foraging bout (**Fig.3B**; species: est=-0.18, z_945_=-2.97, p=0.003). Honey bees also took significantly more foraging bouts to discover all flowers (honey bees: 19 ± 5.7 (SE) foraging bouts; bumblebees: 4 ± 1.2 (SE) foraging bouts) (**Fig.3C**). The flowers that were the farthest from the hive (flowers 2 and 4) were discovered significantly later by honey bees (**Fig.3C;** Wilcoxon test; *F2*: U=22.5, p=0.04; *F4*: U=16, p=0.02; p>0.05 for all other flowers). Moreover, all flowers, except flower 5, were visited in fewer foraging bouts by honey bees than by bumblebees (**Fig.3D;** Wilcoxon test; *F1*: U=71.5, p=0.03; *F2*: U=84, p=0.001; *F3*: U=80.5, p=0.002; *F4*: U=80.5, p=0.004; *F5*: U=66, p=0.08; *F6*: U=76, p=0.01).

**Figure 3:**
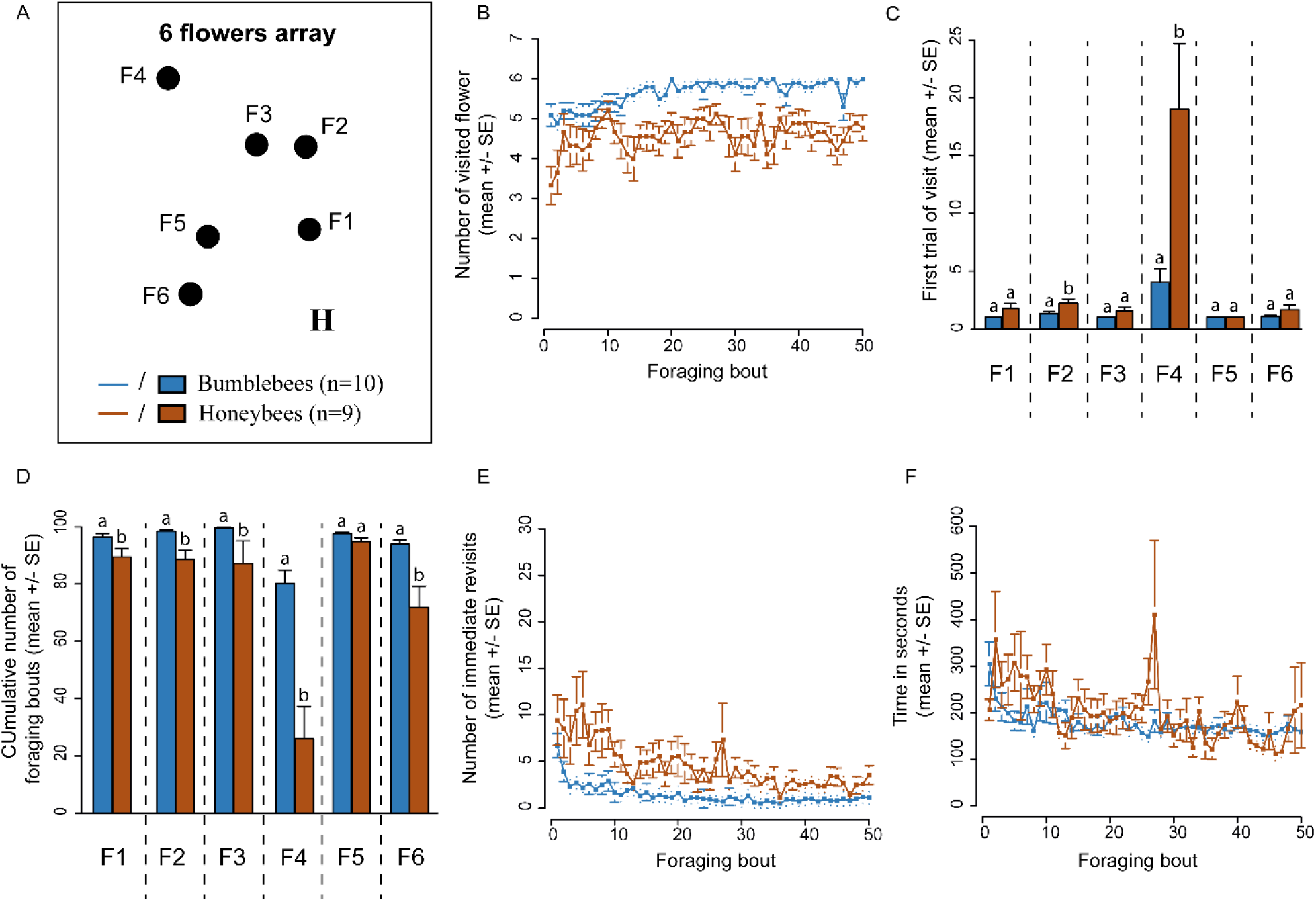
Route formation in the six-flowers array. **(A)** Schematic representation of the flower array. Flowers are labelled F1-F6. H is the colony hive location. **(B)** Number of flowers visited per foraging bouts. **(C)** Cumulated number of foraging bouts before the flower was first visited **(D)** Percentage of foraging bouts for which each flower was visited. **(E) N**umber of immediate revisits to flowers per foraging bout. **(F)** Flight duration per foraging bout. The mean ± standard error is presented for each graph. Different letters indicate a significant difference in **(C)** and **(D)** (Wilcoxon Rank-test for independent data).

Bumblebees and honey bees similarly decreased their number of immediate revisits to flowers with experience (**Fig.3E**; Poisson GLMM; foraging bout: est=-0.03, z_945_=-12.8, p<0.0001; species x foraging bout: est=0.006, z_945_=1.88, p=0.06). Foragers of both species gradually decreased their flight duration (**Fig.3F**; Poisson GLMM; foraging bout: est=-0.006, z_945_=-29.26, p<0.0001). However, once again, honey bees spent significantly more time flying than bumblebees (**Fig.3F**; species: est=0.26, z_945_=2.6, p=0.01; species x foraging bout: est=-0.005, z_945_=-16.63, p<0.0001). Thus overall, honey bees showed a lower increase in route quality with time (Binomial GLMM; foraging bout: est=0.064, z_945_=7.953, p<0.0001; species x foraging bout: est=-0.043, z_945_=-3.087, p=0.002; Fig. 5B).

All bumblebees developed a primary route between the six flowers (n = 10; **Fig.4A**). On average, bumblebees used it in 14 ± 2% (SE) of their foraging bouts, which is higher than expected by chance (binomial test, p<0.0001). The optimal route (F1-F2-F3-F4-F5-F6) was used as primary route by 40% of the bumblebees (4 out of 10 individuals). By contrast, only two honey bees developed a primary route between the six flowers and they used it in 12 ± 2% (SE) of their foraging bouts. The seven remaining honey bees only developed primary routes between five flowers (n=4), four flowers (n=2), or three flowers (n=1; **Fig.4B**). Irrespective of the number of flowers visited, both species showed a similar increase in route similarity (DET) over successive foraging bouts (Binomial GLMM; foraging bout: est=0.265, z_774_=2.817, p=0.005; honey bee x foraging bout: est=4.583e+01, z_774_=0.000, p=1.000; Fig. 5B).

**Figure 4:**
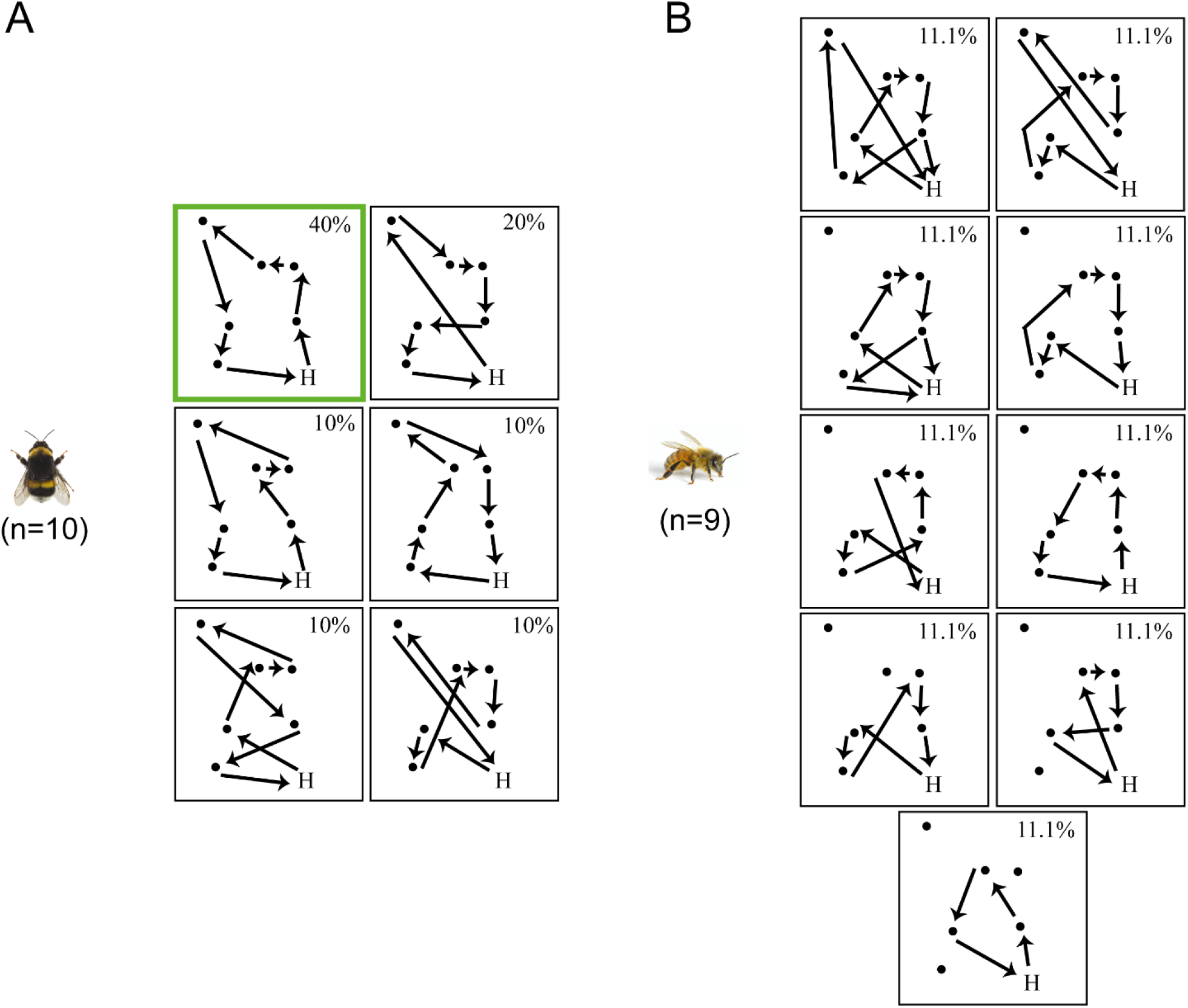
Primary routes on the six-flowers assay. **(A)** Primary route used by bumblebees. **(B)** Primary route used by honeybees **(B)**. Black dots represent flower locations. H is the colony hive location. Arrows indicate the direction in which the bee moved. The percentage indicates the proportion of bees that used this route pattern. Optimal routes are highlighted in green.

**Figure 5.**
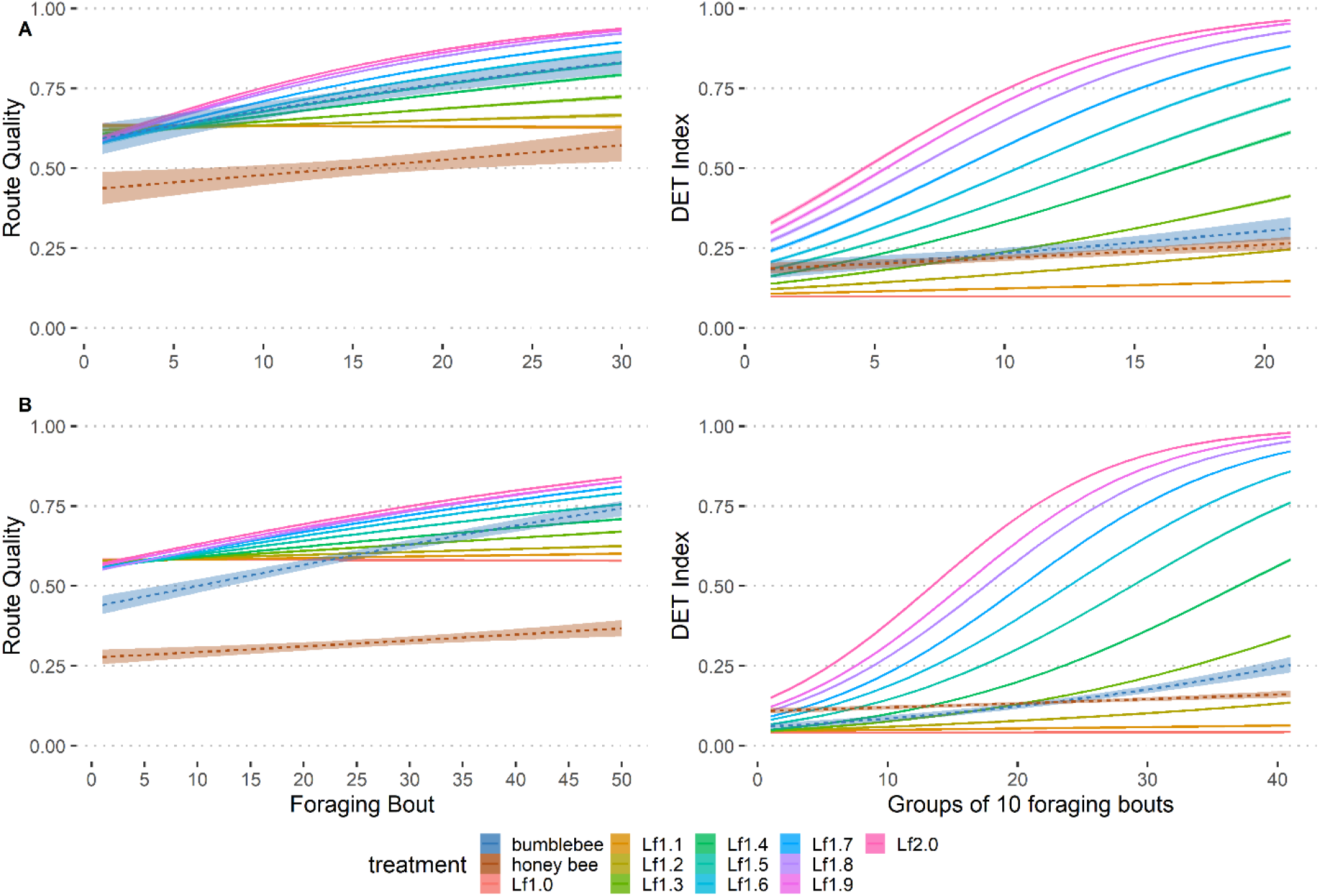
Comparison between observed and simulated data. For each batch of model simulations (named after the value of the learning factor Lf) and both species (bumbleblees and honey bees), the route quality and the route similarity (DET) indices are shown. **A)** Four-flowers array. **B)** Six-flowers array. N = 1000 simulations per Lf. Means are displayed with standard errors.

Thus, foragers of both species increased their foraging efficiency with experience. However, while bumblebees developed a route between the six flowers, honey bees hardly found all flowers and used routes between less flowers.

### Model simulations

To test whether the model of trapline formation proposed in previous studies (Lihoreau et al. 2012b; Reynolds et al. 2013) could explain the observed differences in routing behaviour between bumblebees and honey bees, we compared observed flower visitation sequences to simulated flower visitation sequences, using route quality and route similarity (DET). In each case, we compared the binomial fit of the experimental data to that of the simulations with different values of learning factor (Lf), and rejected all models that showed a significant difference, ultimately providing a set of Lf relevant for each species. The results are summarized in Table 1 (comparisons of slopes) and Table S2 (comparisons of intercepts).

**Table 1:** Statistical comparison between observed and simulated data. Summary of all the p-values obtained when comparing the models to the experimental data, using the route quality and route similarity (DET Index). The p-values displayed are for the slope comparisons. The color code for p-values is ‘green’ <1; ‘yellow’ <0.1; ‘orange’ <0.05; ‘light red’ <0.01; ‘red’ <0.001.

| Array | Variable | Species | Lf1.0 | Lf1.1 | Lf1.2 | Lf1.3 | Lf1.4 | Lf1.5 | Lf1.6 | Lf1.7 | Lf1.8 | Lf1.9 | Lf2.0 |
| --- | --- | --- | --- | --- | --- | --- | --- | --- | --- | --- | --- | --- | --- |
| 4F | Quality | Bumblebee | 0.001 | 0.005 | 0.029 | 0.084 | 0.306 | 0.832 | 0.432 | 0.155 | 0.005 | 9.00e-04 | 2.89e-05 |
| 4F | Quality | Honey bee | 0.014 | 0.070 | 0.054 | 0.577 | 0.332 | 0.070 | 2.92e-04 | 1.13e-06 | 3.42e-12 | 6.77e-12 | 9.70e-14 |
| 4F | Similarity (DET) | Bumblebee | 0.046 | 0.351 | 0.557 | 0.046 | 0.035 | 0.007 | 7.04e-05 | 0.008 | 2.77e-05 | 2.91e-05 | 1.26e-04 |
| 4F | Similarity (DET) | Honey bee | 0.667 | 0.222 | 0.324 | 0.029 | 0.001 | 0.003 | 0.001 | 0.002 | 4.35e-06 | 1.25e-05 | 3.52e-05 |
| 6F | Quality | Bumblebee | 1.42e-06 | 2.05e-07 | 1.20e-06 | 1.22e-04 | 8.72e-04 | 0.154 | 0.743 | 0.154 | 0.018 | 0.059 | 6.78e-05 |
| 6F | Quality | Honey bee | 8.59e-04 | 0.001 | 0.003 | 0.084 | 0.525 | 0.106 | 0.012 | 3.81e-08 | 4.03e-08 | 2.21e-06 | 2.25e-07 |
| 6F | Similarity (DET) | Bumblebee | 0.007 | 0.339 | 0.978 | 0.064 | 0.052 | 0.343 | 0.225 | 0.040 | 0.077 | 0.033 | 0.064 |
| 6F | Similarity (DET) | Honey bee | 0.077 | 0.453 | 0.800 | 0.077 | 0.019 | 0.091 | 0.005 | 9.69e-04 | 3.18e-04 | 0.003 | 6.16e-04 |

We first compared observational and simulated data in the four-flowers array. For bumblebees, the increase of route quality was replicated with Lf ranging from 1.3 to 1.7 (**Fig.5A**; Table 1) and the increase in route similarity (DET) with Lf ranging from 1.1 to 1.2 (**Fig.5B**; Table 1). For honey bees, the increase of route quality was replicated with Ln ranging from 1.1 to 1.5 (**Fig.5A**; Table 1) and the increase of route similarity with Ln ranging from 1.0 to 1.2 (**Fig.5B**; Table 1). Results followed a similar pattern in the six-flowers array. For bumblebees, the increase of route quality was replicated with Lf 1.5, 1.6, 1.7 and 1.9 (**Fig.5C**; Table 1) and the increase of route similarity with Lf ranging from 1.1 to 1.6, and 1.8 and 2.0. (**Fig.5D**; Table 1). For honey bees, route quality was replicated with a Lf ranging from 1.3 to 1.5 (**Fig.5C**; Table 1) and route similarity with a Lf ranging from 1.0 to 1.3 and 1.5 (**Fig.5D**; Table 1). Similar observations were made by comparing the intercepts of experimental data and models (Table S2). Thus overall, the behaviour of honey bees was replicated with lower values of lf than that of bumblebees in both arrays of flowers. This indicates that honey bees were generally slower to develop a route and that their routes were also less efficient.

## Discussion

We compared the routing behaviour of two naturally co-occurring generalist pollinators foraging in the same arrays of artificial flowers. While bumblebees and honey bees established routes between four flowers, only bumblebees did so between six flowers. In these conditions, honey bees took a longer time to discover all the flowers and never integrated them all in a single stable visitation sequence.

Many animals develop foraging routes (or traplines) to efficiently exploit multiple familiar feeding sites that replenish over time (Janzen 1971; Thomson et al. 1997; Buatois and Lihoreau 2016; Gilbert and Singer 1975; Tello-Ramos et al. 2015; Lemke 1984; Wooller et al 1999). Most experiments on trapline foraging have been designed to test whether animals could optimise overall travel distances and to explore the cognitive mechanisms involved in route formation (Ohashi and Thomson 2009; Lihoreau et al. 2013). However, comparing these behaviours across individuals and species can bring important insights into the evolution of spatial cognition and its impact on pollination. Recently, Klein et al. (2017) compared the spatial patterns of bumblebee foragers in different arrays of flowers, showing consistent inter-individual variations in speed and efficiency for route development. Here we adopted a similar strategy to compare the routing behaviour of bumblebees and honey bees, and demonstrated that honey bees are generally slower in developing routes, and that their routes are less efficient than those of bumblebees. This result is consistent with indirect comparisons of semi field data (Pasquaretta et al. 2017). Importantly, this key behavioural difference between honey bees and bumblebees was well-captured by changing a single parameter value (the learning factor) in simulations of a previously proposed model of trapline formation (Lihoreau et al. 2012b, Reynolds et al. 2013). Overall, the behaviour of honey bees was best-replicated with lower learning factors than that of bumblebees.

Why do honey bees and bumblebees do not behave the same? The two species greatly differ in social organization and it is possible that their behavioural strategies are influenced by social constraints. Honey bees use the waggle dance to recruit foragers on profitable sites (von Frisch 1967) and may thus invest less in individual sampling and learning than bumblebees that do not exhibit dance communication (Dornhaus and Chittka 1999). At the collective level, unperfect traplining and dance communication may be more efficient in tropical environments dominated by densely aggregated resources such as blooming trees where honey bees are expected to originate (Dyer and Seeley 1989), whereas perfect traplining may be an efficienct strategy to maximize collective foraging in environments dominated by spatially distributed resources such as small flower patches. The size of colonies are also dramatically different in the two species. While a honey bee colony can contain thousands of workers, a bumblebee colony only contains hundreds. The foraging workforce of honey bees is therefore highly superior, resulting in a potentially more intense activity of exploration to find new resources and exploit them rapidly, than individual learning to monopolize specific resources for long period of times. Recent studies indicate that honey bee foragers are less accurate than bumblebee foragers in many aspects of foraging, for instance when selecting pollen of different qualities (Leonhard and Bluthgen 2012) or searching for a target in an arena (Morawetz and Spaethe 2012). Our results are consistent with these observations since, even in the most complex array of six flowers, bumblebees managed to find every flower, while honey bees never visited them all during the same foraging bout.

These different spatial behaviours between honey bees and bumblebees may also result from differences in bio-energetics. In addition to an obvious difference of body size, foragers of the two species have a very different nectar crop capacities. While it is well accepted that the usual crop capacity of honey bees is about 60 µL (Nunez 1966), the crop of bumblebees is typically around 120 µL (Lihoreau et al. 2010,2012a,b). In contrast to bumblebees that often return home with a full crop, honey bees often abandon non-depleting food sources with a partially filled crop (Schmid-Hempel et al. 1985). This behaviour does not maximize the net rate of energy extraction from the food sources but appears to maximize energetic efficiency (net energetic gain/unit energy expenditure) (Schmid-Hempel et al. 1985). This may explain why honey bees in the six-flowers array rarely visited all flowers during a given foraging bout. Note however that honey bees tested both on the four- and six-flowers arrays did significantly more immediate revisits to empty flowers than bumblebees, as well as flew significantly longer, suggesting that they were searching for additional food. It remains relatively unclear why bumblebees have a crop capacity way more stable than honeybees during a foraging flight. Different studies have shown that metabolic rates tested in bumblebees and honey bees during flight were not so different (Kammer and Heinrich 1974, Heinrich 1975a,b, Schaffer et al. 1979, Harrison and Fewell 2002, Darveau et al. 2014). Consequently, the flight energy could not be an explanation for the differences observed during our experiments. In addition to potential physical constraints, it is more likely that the difference regarding the crop capacity is triggered by the difference in colony size, as well as the number of recruited foragers for both species. Indeed, the low number of foragers in bumblebees, as well as a poor communication about food, have to be balanced by an optimal individual foraging and therefore a fully filled crop to ensure a maximal transport of ressources. However, in honeybees, with a very large number of foragers, as well as a very sophisticated system of recruitment, the individual foraging does not have to be as efficient since other bees will come to help foraging the resources.

The morphological differences between the two species are also accompanied with size differences in sensory organs and brains that could influence routing behaviour. For instance, the bumblebee eye surface is 2 times larger (Streinzer et al. 2013, Streinzer and Spaethe 2014) and its resolution is about 25% better than that of the honey bee eye (Macuda et al. 2001). Consequently, bumblebees are more likely to identify a small object at a long distance. In our experiments, although artificial flowers were designed to be detectable by both bumblebees and honey bees, differences in visual acuity may explain why bumblebees were more efficient at finding all flowers. Beyond visual perception, honey bees and bumblebees can notoriously perform a rich diversity of elemental and non-elemental learning tasks (Giurfa 2013, Perry et al. 2017). Despite these elaborated cognitive abilities in both species, the different body size of the two insects also comes with an obvious difference of brain size (Mares et al. 2005). Although it is still quite debated (Chittka and Niven 2009, Lihoreau et al. 2012c), some studies conducted in different species has shown that a larger brains could be associated with better cognitive performances (Buechel et al. 2018; Herculano-Houzel 2017). For instance, when tested on the same learning task, large-brained females guppy performed better than the small-brain females (Kotrschal et al. 2013). Such observations have also been performed in bees and allowed to highlight correlation between brain size and some specific behaviour. Sayol et al. (2020) showed an association between larger brain could be associated with ecologically specialized bee species when compared with generalist or multi-generation species. More recently, this question has been explored by comparing 32 different bee species brain size after a colour discrimination task. Regardless of the species, bees with a larger brain were more likely to learn the association between the reward and the colour (Collado et al. 2020). It is therefore reasonable to hypothesise that with a larger brain, bumblebees would be cognitively more efficient, explaining their best performance in our experiments.

Our study focused on route development by bees at small spatial scales in the lab. Although bees can forage in these conditions, route optimization is faster and more pronounced at larger spatial scales, where feeding sites are visually isolated from each other and the cost of flying long suboptimal routes is magnified (Lihoreau et al. 2012b, Buatois and Lihoreau 2016; Woodgate et al. 2017). Future studies should therefore explore whether the differences between bumblebees and honey bees are maintained at larger spatial scales. Honey bees and bumblebees can fly to up to ca. 10 kms (Goulson and Stout 2001; Pahl et al. 2011). In the field, the difference in relative energetic cost of flight in the two species, due body size differences, may lead honey bees to invest more in route optimization between distant feeding sites. Ultimately, a more systematic analysis and comparisons of the spatial behaviours of key pollinators will help clarify their impact on pollen transfer and potential complementarity or redundancy for plant reproduction and crop yield (Garibaldi et al. 2016).

## Acknowledgements

This work was funded by the CNRS and three grants from the Agence Nationale de la Recherche to ML (ANR-16-CE02-0002-01; ANR-19-CE37-0024; ANR-20-ERC8-0004-01). While writing, AB was funded by Fyssen. TD was funded by a co-tutelle PhD grant of the University Paul Sabatier (Toulouse) and Macquarie University (Sydney).

