## Supplementary for "Bumblebees develop more efficient traplines than honey bees"

**Text S1. R code used to generate arrays of flowers.**

**#### Code to generate all route combination possible (example with 6 flowers) ####**

- seq_flow = c(1,2,3,4,5,6) # Define a 6 flowers sequence
- mat = expand.grid (seq_flow, seq_flow, seq_flow, seq_flow, seq_flow, seq_flow, KEEP.OUT.ATTRS = F) # provide all 6 flowers sequences possible
- line_to_remove=NULL #create a NULL vector

- for (i in 1: nrow(mat)) # Remove the repetition inside the provided sequences

  { if(length(which(duplicated(t(mat[i,]))==T))>0) line_to_remove=c(line_to_remove,i) }

- mat<-mat[-line_to_remove,] # New matrix without repetition
- mat<-cbind(0,mat,0) # add a 0 at the beginning and at the end of each sequence
- rownames(mat)<-1:nrow(mat) #rename the lines

**#### Generate random conformation ####**

- library(spatstat) #package needed

#Matrix B corresponds to X coordinates matrix C to Y coordinates

- B=mat
- C=mat

### generate 7 points randomly on the surface of the room #

- Flow=runifpoint(7,win=owin(c(0,470),c(0,670))) # generate 7 point randomly on the surface
- X=Flow$x #Define vector X as containing X coordinates generate by runifpoint.
- Y=Flow$y #Define vector Y as containing Y coordinates generate by runifpoint.
- X[1]=400 #Define X Hive coordinate as fixed at 400 cm
- Y[1]=50 #Define Y Hive coordinates as fixed at 50 cm

**##### Calculate the maximal distance, as well as the shorter distance to visit straightly each point of the conformation previously obtained ####**

- for (i in c(0:6)) # replace Flower number by its X coordinates in B et Y coordinates in C

{  B[B==i]<-X[i+1]  C[C==i]<-Y[i+1] }

- mat_distance = NULL #create a NULL vector to store the calculated distance
- for (i in 1:nrow(mat)) #calcule la distance totale pour chaque combinaison

  {

  Distance=sqrt(((B[i,2])-(B[i,1]))^2+((C[i,2])-(C[i,1]))^2)+sqrt(((B[i,3])(B[i,2]))^2+((C[i,3])-(C[i,2]))^2)+sqrt(((B[i,4])(B[i,3]))^2+((C[i,4])-(C[i,3]))^2)+sqrt(((B[i,5])(B[i,4]))^2+((C[i,5])-(C[i,4]))^2)+sqrt(((B[i,6])(B[i,5]))^2+((C[i,6])(C[i,5]))^2)+sqrt(((B[i,7])-(B[i,6]))^2+((C[i,7])-(C[i,6]))^2)+sqrt(((B[i,8])(B[i,7]))^2+((C[i,8])-(C[i,7]))^2)

  mat_distance=c(mat_distance,Distance) }

- mat3=cbind(mat,distance=mat_distance) #create a matrix with every route and the corresponding distance

- minseg=mat3[which((mat3$distance)==min(mat3$distance)),] #Highlight the route with the minimal distance

- maxseg=mat3[which((mat3$distance)==max(mat3$distance)),] #Highlight the route with the maximal distance

**#### Calculate the distance using the closest neighbor visits strategy####**

- seq_flow<-c(1:6) #Define a sequence of flowers going from 1 to 6
- XY_flow<-cbind(x=X[-1],y=Y[-1],flower=seq_flow) # Coordinates Matrix
- mat_nn<-NULL
- mat_voisin<-(NULL)

- for (i in 1:nrow(XY_flow)) #Minimal distance between the hive and the first flower

  { DistanceVoisin = sqrt((((X[1])-(XY_flow[i,1]))^2)+(((Y[1])-(XY_flow[i,2]))^2)) mat_voisin=rbind(mat_voisin,c(DistanceVoisin,i))
  min_dist_nn<-which((mat_voisin[,1])==min(mat_voisin[,1])) }

- mat_nn=rbind(mat_nn, mat_voisin[min_dist_nn,])

- XY_flow<-XY_flow[-which(XY_flow[,3]==mat_nn[nrow(mat_nn),2]),] #remove the flower that is closer to the hive

- XY_flow<-as.data.frame(XY_flow)

- for (j in 1: 5) {

  mat_voisin=NULL
  for (i in 1:nrow(XY_flow)) # Minimal distance between the flower that are closer to each other

  {

  x0<-X[mat_nn[nrow(mat_nn),2]+1]
  y0<-Y[mat_nn[nrow(mat_nn),2]+1]
  DistanceVoisin = sqrt((x0-(XY_flow[i,1]))^2+(y0-(XY_flow[i,2]))^2) mat_voisin=rbind(mat_voisin,c(DistanceVoisin,XY_flow[i,3])) # Distance for each conformation
  }

  min_dist_nn<-which((mat_voisin[,1])==min(mat_voisin[,1]))
  mat_nn=rbind(mat_nn,mat_voisin[min_dist_nn,])

XY_flow<-XY_flow[-which(XY_flow[,3]==mat_nn[nrow(mat_nn),2]),] #Remove the closest flower from the hive }

**#The minimal distance between the last visited flower and the hive**

- dist_last_flow_nest<-sqrt((X[1]-(X[mat_nn[6,2]]))^2+ (Y[1]-(Y[mat_nn[6,2]]))^2)
- mat_nn=rbind(mat_nn, c(dist_last_flow_nest,0))
- seq_nn<-c(0,mat_nn[,2])
- D=seq_nn
- E=seq_nn

- for (i in c(0:6)) # replace flower number by its X coordinate {

D[D==i]<-X[i+1] E[E==i]<-Y[i+1] }

- Distance_nn = sqrt(((D[2])-(D[1]))^2+((E[2])-(E[1]))^2) + sqrt(((D[3])-(D[2]))^2+((E[3])(E[2]))^2) + sqrt(((D[4])-(D[3]))^2+((E[4])-(E[3]))^2) + sqrt(((D[5])-(D[4]))^2+((E[5])(E[4]))^2) + sqrt(((D[6])-(D[5]))^2+((E[6])-(E[5]))^2) + sqrt(((D[7])-(D[6]))^2+((E[7])(E[6]))^2) + sqrt(((D[8])-(D[7]))^2+((E[8])-(E[7]))^2)
- Dist_nn=c(seq_nn,Distance_nn)

- if(minseg$distance[1]!= Distance_nn) # If minimal distance and closest neighbour distance are different, the distance informations, as well as the coordinates of the conformation are saved.

  { print("DIFFERENCE") write.csv2(rbind(minseg,maxseg,Dist_nn),file="Distance.csv") write.csv2(rbind(X,Y),file="coordoninates.csv") }

Once several conformations have been saved, the ratio between minimal distance and closest neighbour distance was calculated. Only the conformation with the higher ratio has been chosen by the end.

#### Table S1

**Table S1: Four-flower visitation sequences (excluding revisits) for each bumblebee tested in the four-flowers array.** Numbers (1-4) in tables refer to unique flowers (see details in Fig. S1), and colour codes refer to a unique flower sequence. Incomplete sequences (not included in the analyses of sequence repeatability) are in white. Sequences in columns are sorted in chronological order (from foraging bout 1 to foraging bout 30).

| | Bee 1 | Bee 2 | Bee 3 | Bee 4 | Bee 5 | Bee 6 | Bee 7 | Bee 8 | Bee 9 | Bee 10 |
|---|---|---|---|---|---|---|---|---|---|---|
| **Bout1** | 413 | 4132 | 1423 | 14 | 431 | 412 | 41 | 4132 | 143 | 143 |
| **Bout2** | 3214 | 4123 | 143 | 413 | 4312 | 134 | 143 | 134 | 413 | 413 |
| **Bout3** | 2341 | 4321 | 143 | 431 | 3142 | 314 | 1432 | 413 | 413 | 134 |
| **Bout4** | 2341 | 4321 | 143 | 413 | 3421 | 3142 | 1432 | 413 | 413 | 413 |
| **Bout5** | 3412 | 1432 | 1432 | 431 | 413 | 3214 | 2314 | 4123 | 4132 | 431 |
| **Bout6** | 1234 | 3214 | 1432 | 431 | 412 | 142 | 4132 | 413 | 1342 | 4132 |
| **Bout7** | 1234 | 4132 | 4321 | 132 | 4312 | 3214 | 4132 | 4132 | 413 | 4132 |
| **Bout8** | 3412 | 3241 | 1342 | 431 | 3214 | 1432 | 4231 | 4321 | 4321 | 4132 |
| **Bout9** | 1234 | 3241 | 1423 | 413 | 4321 | 1234 | 4321 | 4123 | 4132 | 4123 |
| **Bout10** | 324 | 3214 | 1432 | 4123 | 4132 | 1423 | 1243 | 4321 | 4132 | 2314 |
| **Bout11** | 1234 | 1234 | 1342 | 413 | 4321 | 134 | 4132 | 4321 | 4132 | 4123 |
| **Bout12** | 1234 | 2134 | 4321 | 4132 | 432 | 3214 | 413 | 432 | 4132 | 412 |
| **Bout13** | 1234 | 3214 | 1432 | 4132 | 3214 | 3214 | 1234 | 4132 | 4132 | 4123 |
| **Bout14** | 1234 | 4123 | 4321 | 4123 | 3241 | 1324 | 4123 | 321 | 4132 | 4123 |
| **Bout15** | 1234 | 4123 | 1342 | 413 | 2134 | 1324 | 4321 | 4321 | 4132 | 4123 |
| **Bout16** | 1423 | 3241 | 4321 | 4123 | 1432 | 3214 | 4132 | 4321 | 4132 | 1234 |
| **Bout17** | 2314 | 3241 | 4132 | 4132 | 1324 | 1432 | 4132 | 4321 | 4132 | 4132 |
| **Bout18** | 1234 | 3214 | 4312 | 4132 | 312 | 4321 | 4132 | 4321 | 432 | 4123 |
| **Bout19** | 1234 | 4312 | 4312 | 4132 | 432 | 132 | 123 | 4132 | 4321 | 1234 |
| **Bout20** | 1234 | 4321 | 1324 | 4132 | 3214 | 4321 | 4321 | 4321 | 4132 | 1234 |
| **Bout21** | 2341 | 4312 | 4321 | 4132 | 4312 | 1432 | 4132 | 4321 | 4132 | 4123 |
| **Bout22** | 2341 | 4312 | 1342 | 4132 | 4312 | 3214 | 4132 | 4321 | 4132 | 1234 |
| **Bout23** | 1234 | 3241 | 4321 | 413 | 4321 | 1324 | 4132 | 4321 | 4321 | 3124 |
| **Bout24** | 2341 | 3241 | 3214 | 4132 | 3214 | 412 | 4312 | 4321 | 4132 | 1234 |
| **Bout25** | 1234 | 3214 | 1324 | 4132 | 3214 | 1324 | 4321 | 4321 | 4132 | 4132 |
| **Bout26** | 2341 | 142 | 4321 | 3214 | 314 | 4132 | 4123 | 3214 | 4132 | 4123 |
| **Bout27** | 2341 | 3214 | 4321 | 412 | 321 | 1324 | 132 | 4321 | 4321 | 4123 |
| **Bout28** | 2341 | 3241 | 4321 | 4132 | 3214 | 4321 | 4321 | 4321 | 1324 | 2341 |
| **Bout29** | 2341 | 4123 | 4321 | 4132 | 3214 | 1324 | 4231 | 4321 | 4321 | 4123 |
| **Bout30** | 1234 | 4321 | 4321 | 4123 | 3214 | 132 | 412 | 4321 | 4321 | 4123 |

### Table S2

**Table S2: Four-flower visitation sequences (excluding revisits) for each honey bee tested in the four-flowers array.** Numbers (1-4) in tables refer to unique flowers (see details in Fig. S1), and colour codes refer to a unique flower sequence. Incomplete sequences (not included in the analyses of sequence repeatability) are in white. Sequences in columns are sorted in chronological order (from foraging bout 1 to foraging bout 30).

| | Bee 1 | Bee 2 | Bee 3 | Bee 4 | Bee 5 | Bee 6 | Bee 7 | Bee 8 | Bee 9 | Bee 10 |
|---|---|---|---|---|---|---|---|---|---|---|
| **Bout 1** | 41 | 14 | 341 | 41 | 4312 | 14 | 41 | 143 | 41 | 41 |
| **Bout 2** | 41 | 14 | 41 | 41 | 4123 | 14 | 41 | 14 | 41 | 4 |
| **Bout 3** | 4132 | 4123 | 14 | 4123 | 4132 | 143 | 41 | 143 | 41 | 14 |
| **Bout 4** | 143 | 1234 | 143 | 143 | 1432 | 123 | 41 | 434 | 41 | 4132 |
| **Bout 5** | 123 | 1432 | 4321 | 432 | 4321 | 3214 | 41 | 23 | 1423 | 3214 |
| **Bout 6** | 1432 | 1234 | 1432 | 1 | 1324 | 3213 | 413 | 134 | 1 | 4321 |
| **Bout 7** | 412 | 123 | 4321 | 4321 | 1234 | 1234 | 4132 | 413 | 1423 | 4321 |
| **Bout 8** | 1234 | 12 | 4123 | 3214 | 3421 | 321 | 4132 | 1234 | 1432 | 4321 |
| **Bout 9** | 1234 | 12 | 4321 | 4321 | 1423 | 1234 | 1243 | 1324 | 4123 | 4321 |
| **Bout 10** | 4123 | 123 | 4132 | 342 | 3421 | 123 | 4321 | 1432 | 1234 | 4321 |
| **Bout 11** | 123 | 234 | 4132 | 324 | 3421 | 3124 | 1432 | 1432 | 1324 | 4321 |
| **Bout 12** | 123 | 231 | 3142 | 4321 | 1234 | 123 | 4321 | 1234 | 123 | 2134 |
| **Bout 13** | 1234 | 2341 | 4132 | 321 | 4231 | 1234 | 4321 | 1234 | 1 | 3214 |
| **Bout 14** | 4123 | 123 | 1432 | 2314 | 1234 | 1234 | 4321 | 1234 | 1234 | 3214 |
| **Bout 15** | 4123 | 2314 | 1234 | 3214 | 1432 | 3214 | 4321 | 1234 | 1234 | 3214 |
| **Bout 16** | 4123 | 231 | 4123 | 4321 | 3124 | 231 | 4321 | 1234 | 231 | 2314 |
| **Bout 17** | 2314 | 23 | 4123 | 4321 | 1234 | 123 | 4321 | 1234 | 1234 | 3214 |
| **Bout 18** | 123 | 1423 | 2314 | 3124 | 3214 | 231 | 1234 | 1234 | 2341 | 2143 |
| **Bout 19** | 1234 | 2314 | 2314 | 431 | 1234 | 1234 | 4321 | 1234 | 1234 | 3241 |
| **Bout 20** | 4123 | 432 | 4132 | 3124 | 4321 | 123 | 4321 | 1234 | 2341 | 3214 |
| **Bout 21** | 4123 | 123 | 4123 | 312 | 3241 | 1234 | 4321 | 1234 | 4123 | 3214 |
| **Bout 22** | 1234 | 4123 | 3214 | 3142 | 23 | 1234 | 4321 | 1324 | 132 | 2314 |
| **Bout 23** | 4123 | 1423 | 2314 | 3124 | 1234 | 1234 | 4321 | 1234 | 2341 | 3214 |
| **Bout 24** | 2314 | 123 | 1234 | 3124 | 23 | 1234 | 4321 | 1234 | 23 | 3241 |
| **Bout 25** | 4123 | 1234 | 4123 | 4321 | 1324 | 1234 | 4321 | 1342 | 2341 | 4321 |
| **Bout 26** | 4123 | 4132 | 4123 | 3142 | 1342 | 1234 | 3214 | 1234 | 1234 | 4321 |
| **Bout 27** | 14 | 2314 | 1234 | 2134 | 1342 | 123 | 4321 | 1234 | 1234 | 2314 |
| **Bout 28** | 4123 | 1234 | 1234 | 4321 | 231 | 1234 | 4321 | 1234 | 1234 | 2314 |
| **Bout 29** | 1234 | 1234 | 1234 | 2143 | 2 | 123 | 4321 | 1234 | 1342 | 3214 |
| **Bout 30** | 1234 | 1234 | 1234 | 3421 | 3421 | 1234 | 4321 | 1234 | 23 | 3214 |

#### Table S3

**Table S3: Six-flower visitation sequences (excluding revisits) for each bumblebee tested in the six-flowers array.** Numbers (1-6) in tables refer to unique flowers (see details in Fig. S1), and colour codes refer to a unique flower sequence. Incomplete sequences (not included in the analyses of sequence repeatability) are in white. Sequences in columns are sorted in chronological order (from foraging bout 1 to foraging bout 50).

| | Bee 1 | Bee 2 | Bee 3 | Bee 4 | Bee 5 | Bee 6 | Bee 7 | Bee 8 | Bee 9 | Bee 10 |
|---|---|---|---|---|---|---|---|---|---|---|
| **Bout1** | 35612 | 35612 | 56132 | 153264 | 456132 | 5316 | 531642 | 1532 | 351642 | 5613 |
| **Bout2** | 43216 | 35216 | 23156 | 132546 | 23561 | 5123 | 35612 | 53612 | 13526 | 5316 |
| **Bout3** | 21563 | 35162 | 3524 | 125364 | 53612 | 65123 | 653124 | 65132 | 563214 | 52312 |
| **Bout4** | 23165 | 345126 | 45613 | 123456 | 65321 | 32165 | 536142 | 52316 | 32561 | 5321 |
| **Bout5** | 45632 | 53216 | 45613 | 123456 | 13526 | 2136 | 435216 | 53612 | 53162 | 53216 |
| **Bout6** | 312456 | 32156 | 234 | 123456 | 63215 | 64312 | 153264 | 15632 | 53216 | 51236 |
| **Bout7** | 16532 | 65321 | 251346 | 123456 | 65321 | 562 | 43216 | 15632 | 356142 | 53216 |
| **Bout8** | 324561 | 23156 | 1235 | 123456 | 53261 | 32561 | 213564 | 1253 | 356124 | 65321 |
| **Bout9** | 456132 | 32156 | 23561 | 123456 | 15632 | 21563 | 432156 | 53216 | 324516 | 32165 |
| **Bout10** | 532416 | 56132 | 3245 | 124536 | 65342 | 46512 | 651234 | 153264 | 561234 | 56321 |
| **Bout11** | 123456 | 321564 | 23456 | 123456 | 6532 | 34215 | 43215 | 631254 | 563214 | 63245 |
| **Bout12** | 453612 | 51236 | 1234 | 123456 | 56132 | 34521 | 123546 | 12563 | 531624 | 32456 |
| **Bout13** | 453621 | 56123 | 563214 | 234516 | 352164 | 21563 | 345124 | 53124 | 352164 | 32156 |
| **Bout14** | 123465 | 43156 | 23456 | 213456 | 653214 | 53214 | 432165 | 563124 | 324561 | 32156 |
| **Bout15** | 123546 | 32156 | 432156 | 234156 | 12356 | 512634 | 653214 | 563214 | 563214 | 53216 |
| **Bout16** | 125634 | 432156 | 234156 | 234561 | 65324 | 653412 | 651234 | 321456 | 563214 | 65321 |
| **Bout17** | 234561 | 561234 | 23516 | 234561 | 132456 | 53216 | 432156 | 124356 | 563124 | 532146 |
| **Bout18** | 563142 | 32156 | 53216 | 234561 | 15324 | 12356 | 653214 | 12356 | 563214 | 561234 |
| **Bout19** | 234156 | 65132 | 21345 | 234561 | 12356 | 321564 | 653214 | 12356 | 653214 | 653214 |
| **Bout20** | 321564 | 432156 | 134265 | 234561 | 156324 | 632154 | 512364 | 125364 | 653142 | 563214 |
| **Bout21** | 123654 | 432156 | 421563 | 234516 | 15634 | 43215 | 321546 | 153264 | 345216 | 53261 |
| **Bout22** | 32451 | 653214 | 534216 | 234561 | 153246 | 321564 | 561234 | 123564 | 651234 | 53216 |
| **Bout23** | 651234 | 321654 | 12356 | 234461 | 51234 | 651234 | 653421 | 563412 | 651234 | 653241 |
| **Bout24** | 123456 | 432156 | 234165 | 324156 | 153246 | 12356 | 651234 | 234516 | 324561 | 532146 |
| **Bout25** | 132564 | 316524 | 65312 | 324561 | 234516 | 345621 | 651234 | 123564 | 345612 | 5321 |
| **Bout26** | 235614 | 563214 | 123456 | 356124 | 132456 | 651234 | 432156 | 123456 | 356214 | 65321 |
| **Bout27** | 126534 | 324156 | 534216 | 324561 | 334251 | 653214 | 512346 | 12356 | 324561 | 532146 |
| **Bout28** | 123456 | 564321 | 12356 | 324516 | 651324 | 532164 | 651234 | 64321 | 653421 | 542316 |
| **Bout29** | 123456 | 653214 | 521634 | 325614 | 132546 | 623451 | 654321 | 124356 | 563142 | 53261 |
| **Bout30** | 561324 | 213456 | 123546 | 324561 | 123564 | 532641 | 654321 | 123456 | 132564 | 53264 |
| **Bout31** | 512346 | 532146 | 123465 | 356124 | 532461 | 321546 | 51234 | 561234 | 341562 | 53216 |
| **Bout32** | 456312 | 213654 | 234561 | 321654 | 324165 | 532614 | 653214 | 132456 | 342156 | 5321 |
| **Bout33** | 352416 | 432165 | 21653 | 345612 | 532164 | 653241 | 432156 | 132564 | 345612 | 65321 |
| **Bout34** | 435612 | 564321 | 234561 | 316425 | 132456 | 321564 | 321564 | 134256 | 653214 | 532416 |
| **Bout35** | 465123 | 234561 | 12365 | 342561 | 234561 | 321546 | 432156 | 123456 | 563412 | 534216 |
| **Bout36** | 321546 | 432156 | 123564 | 345612 | 653124 | 653214 | 561234 | 123456 | 563412 | 532641 |
| **Bout37** | 435612 | 456321 | 123465 | 324561 | 16532 | 531246 | 651234 | 12345 | 653214 | 53246 |
| **Bout38** | 123456 | 561234 | 653214 | 345612 | 132456 | 653421 | 561234 | 5326 | 653421 | 5316 |
| **Bout39** | 615342 | 432165 | 653421 | 456321 | 56324 | 653421 | 432156 | 123456 | 345612 | 563241 |
| **Bout40** | 564321 | 453216 | 123564 | 412356 | 132465 | 321456 | 651432 | 12365 | 321645 | 563214 |
| **Bout41** | 123564 | 345621 | 12356 | 456123 | 134256 | 653421 | 432156 | 123456 | 13456 | 532164 |
| **Bout42** | 123546 | 321546 | 123456 | 432156 | 132564 | 534216 | 432165 | 123456 | 23416 | 53241 |
| **Bout43** | 123456 | 324156 | 345612 | 432516 | 653214 | 653421 | 653214 | 12345 | 342156 | 532416 |
| **Bout44** | 123564 | 456321 | 65321 | 423165 | 132564 | 653421 | 432156 | 123456 | 456321 | 321654 |
| **Bout45** | 345216 | 564321 | 234561 | 563124 | 132564 | 651234 | 654321 | 123456 | 432156 | 563241 |
| **Bout46** | 123456 | 34216 | 123456 | 563241 | 132456 | 156342 | 213564 | 123456 | 561234 | 532416 |
| **Bout47** | 432156 | 432 | 12354 | 532461 | 43256 | 653421 | 123654 | 1234 | 432561 | 532164 |
| **Bout48** | 431256 | 532164 | 123456 | 536241 | 563214 | 653241 | 432165 | 123456 | 563214 | 532416 |
| **Bout49** | 15326 | 432156 | 532164 | 532416 | 132456 | 652341 | 651234 | 123456 | 321564 | 532416 |
| **Bout50** | 234561 | 321564 | 123456 | 563214 | 563214 | 653421 | 123564 | 123456 | 312456 | 532461 |

### Table S4

**Table S4: Six-flower visitation sequences (excluding revisits) for each honey bee tested in the six-flowers array.** Numbers (1-6) in tables refer to unique flowers (see details in Fig. S1), and colour codes refer to a unique flower sequence. Incomplete sequences (not included in the analyses of sequence repeatability) are in white. Sequences in columns are sorted in chronological order (from foraging bout 1 to foraging bout 50).

| | Bee 1 | Bee 2 | Bee 3 | Bee 4 | Bee 5 | Bee 6 | Bee 7 | Bee 8 | Bee 9 |
|---|---|---|---|---|---|---|---|---|---|
| Bout1 | 56123 | 15623 | 5 | 5136 | 5632 | 5316 | 51 | 51 | 563 |
| Bout2 | 1613 | 5136 | | 351 | 5362 | 351 | 215364 | 15623 | 5631 |
| Bout3 | 16231 | 13265 | 5 | 35612 | 53261 | 56132 | 51632 | 53216 | 563124 |
| Bout4 | 1 | 56132 | 51324 | 5132 | 513 | 156232 | 564321 | 5123 | 12356 |
| Bout5 | 1 | 56132 | 5631 | 6513 | 32561 | 35216 | 126345 | 5216 | 23516 |
| Bout6 | 1 | 13256 | 5123 | 132564 | 5312 | 6535 | 563214 | 521 | 12536 |
| Bout7 | 1623 | 15632 | 13 | 13524 | 5312 | 5316 | 563214 | 5612 | 12356 |
| Bout8 | 1632 | 5132 | 13256 | 135642 | 5321 | 52316 | 532164 | 5216 | 56123 |
| Bout9 | 1236 | 15362 | 56123 | 123456 | 32156 | 15632 | 532146 | 5612 | 32561 |
| Bout10 | 56123 | 56123 | 532164 | 134562 | 23516 | 56123 | 123564 | 5216 | 53216 |
| Bout11 | 56123 | 53216 | 321564 | 134 | 5321 | 31526 | 53214 | 5612 | 12356 |
| Bout12 | 51236 | 65321 | 231564 | 1534 | 5312 | 561 | 341562 | 56 | 12356 |
| Bout13 | 5612 | 13265 | 1325 | 5134 | 3215 | 5 | 532146 | 5612 | 13256 |
| Bout14 | 56312 | 65321 | 5 | 15 | 35612 | 315 | 563214 | 5126 | 12356 |
| Bout15 | 52563 | 23165 | 5123 | 345 | 32156 | 5632 | 512364 | 5216 | 12356 |
| Bout16 | 52365 | 56321 | 23 | 5342 | 3215 | 56132 | 532164 | 56123 | 12356 |
| Bout17 | 52653 | 65321 | 51234 | 135 | 3215 | 32561 | 534216 | 123 | 12356 |
| Bout18 | 5652 | 123465 | 123 | 135 | 3215 | 32156 | 532146 | 2315 | 12356 |
| Bout19 | 512 | 53216 | 51234 | 31524 | 2156 | 53216 | 321654 | 5621 | 13256 |
| Bout20 | 5532 | 12356 | 132564 | 1534 | 32156 | 12356 | 563214 | 5231 | 13256 |
| Bout21 | 5236 | 13652 | 12345 | 342165 | 231 | 56213 | 56321 | 562 | 12356 |
| Bout22 | 56 | 53216 | 12345 | 53461 | 32156 | 53216 | 564213 | 5612 | 13256 |
| Bout23 | 52356 | 12536 | 324 | 651342 | 3215 | 5321 | 564321 | 5216 | 12356 |
| Bout24 | 51236 | 65321 | 132564 | 32415 | 3215 | 12356 | 532164 | 5612 | 12563 |
| Bout25 | 52653 | 13256 | 532146 | 1325 | 3215 | 356124 | 532164 | 5216 | 12356 |
| Bout26 | 553214 | 123 | 564213 | 314265 | 315 | 13265 | 532146 | 5162 | 13562 |
| Bout27 | 623456 | 53216 | 563214 | 342156 | 3215 | 51326 | 56432 | 5612 | 12563 |
| Bout28 | 63216 | 23156 | 53216 | 34521 | 3561 | 15623 | 532164 | 5216 | 12563 |
| Bout29 | 6 | 31265 | 15324 | 134265 | 3215 | 32165 | 653421 | 5126 | 12356 |
| Bout30 | 325 | 231 | 563142 | 135642 | 3215 | 321 | 532 | 5126 | 12356 |
| Bout31 | 51 | 53216 | 532164 | 134256 | 321 | 53126 | 53216 | 5162 | 12356 |
| Bout32 | 532 | 12365 | 532164 | 134526 | 3215 | 23 | 534126 | 5621 | 12356 |
| Bout33 | 5132 | 65321 | 53 | 13452 | 235 | 12356 | 53421 | 51263 | 12356 |
| Bout34 | 51326 | 32165 | 56324 | 135624 | 3256 | 564231 | 534216 | 6521 | 12365 |
| Bout35 | 3215 | 65321 | 5321 | 12354 | 3215 | 56 | 65312 | 512 | 12356 |
| Bout36 | 5132 | 56321 | 53126 | 354126 | 3256 | 5 | 5321 | 56124 | 23156 |
| Bout37 | 5123 | 53216 | 5134 | 345216 | 3526 | 532146 | 534216 | 51263 | 12356 |
| Bout38 | 5123 | 12 | 51324 | 123546 | 3215 | 13256 | 653214 | 56321 | 12356 |
| Bout39 | 3215 | 12356 | 532164 | 32154 | 3215 | 56321 | 532416 | 6321 | 12356 |
| Bout40 | 5123 | 132 | 532146 | 35126 | 3512 | 56321 | 534216 | 65213 | 12356 |
| Bout41 | 51362 | 65321 | 563214 | 34512 | 3215 | 132 | 536214 | 52146 | 12356 |
| Bout42 | 56123 | 653 | 563214 | 3546 | 321 | 56321 | 563214 | 5621 | 23156 |
| Bout43 | 5321 | 32165 | 534216 | 31456 | 3215 | 6513 | 53216 | 5621 | 12356 |
| Bout44 | 5321 | 13256 | 51236 | 1235 | 32156 | 65132 | 564213 | 6521 | 3561 |
| Bout45 | 5132 | 53216 | 532 | 132 | 3215 | 651324 | 534612 | 6521 | 35621 |
| Bout46 | 5132 | 12356 | 56321 | 325 | 32 | 56321 | 53462 | 5621 | 56123 |
| Bout47 | 3215 | 12536 | 563 | 135 | 32516 | 61235 | 653241 | 5621 | 12365 |
| Bout48 | 5123 | 123 | 532164 | 324516 | 1325 | 56321 | 563421 | 5621 | 12356 |
| Bout49 | 51326 | 12365 | 53214 | 13542 | 3215 | 6513 | 563214 | 56213 | 12356 |
| Bout50 | 5231 | 12356 | 532 | 325146 | 35612 | 56123 | 563214 | 5621 | 12356 |

#### Table S5

**Table S5: Statistical comparison between observed and simulated data.** Summary of all the p-values obtained when comparing the models to the experimental data, using the route quality and route similarity (DET Index). The p-values displayed are for the intercept comparisons. Each colour corresponds to a specific p-value (green: $p>0.1$; yellow: $0.05<p<0.1$; orange: $0.01<p<0.05$; red: $0.001<p<0.01$; brown: $0.001<p$).

| Array | Variable | Species | Ln1.0 | Ln1.1 | Ln1.2 | Ln1.3 | Ln1.4 | Ln1.5 | Ln1.6 | Ln1.7 | Ln1.8 | Ln1.9 | Ln2.0 |
|---|---|---|---|---|---|---|---|---|---|---|---|---|---|
| 4F | Quality | Bumblebee | 0.115 | 0.334 | 0.650 | 0.618 | 0.578 | 0.900 | 0.892 | 0.903 | 0.666 | 0.734 | 0.547 |
| 4F | Quality | Honeybee | 3.98e-08 | 1.02e-05 | 1.14e-06 | 2.41e-05 | 1.34e-04 | 1.62e-04 | 0.004 | 0.008 | 0.014 | 0.001 | 5.92e-04 |
| 4F | Similarity (DET) | Bumblebee | 0.05 | 8.25e-06 | 0.334 | 0.108 | 0.957 | 0.893 | 0.929 | 0.086 | 0.181 | 0.065 | 0.018 |
| 4F | Similarity (DET) | Honeybee | 6.79e-08 | 5.77e-04 | 0.021 | 0.263 | 0.794 | 0.374 | 0.108 | 0.009 | 0.020 | 0.002 | 3.76e-04 |
| 6F | Quality | Bumblebee | 1.00e-04 | 7.85e-06 | 2.89e-05 | 2.98e-04 | 3.11e-04 | 0.006 | 0.075 | 0.045 | 0.057 | 0.004 | 0.060 |
| 6F | Quality | Honeybee | 2.00e-16 | 2.00e-16 | 2.00e-16 | 2.00e-16 | 4.39e-16 | 2.23e-12 | 3.04e-13 | 1.52e-09 | 2.88e-11 | 2.00e-16 | 2.00e-16 |
| 6F | Similarity (DET) | Bumblebee | 0.645 | 0.348 | 0.139 | 0.108 | 0.480 | 0.325 | 0.226 | 0.301 | 0.107 | 0.104 | 0.035 |
| 6F | Similarity (DET) | Honeybee | 0.494 | 0.006 | 0.168 | 0.310 | 0.521 | 0.448 | 0.754 | 0.681 | 0.505 | 0.183 | 0.151 |

#### Figure S1

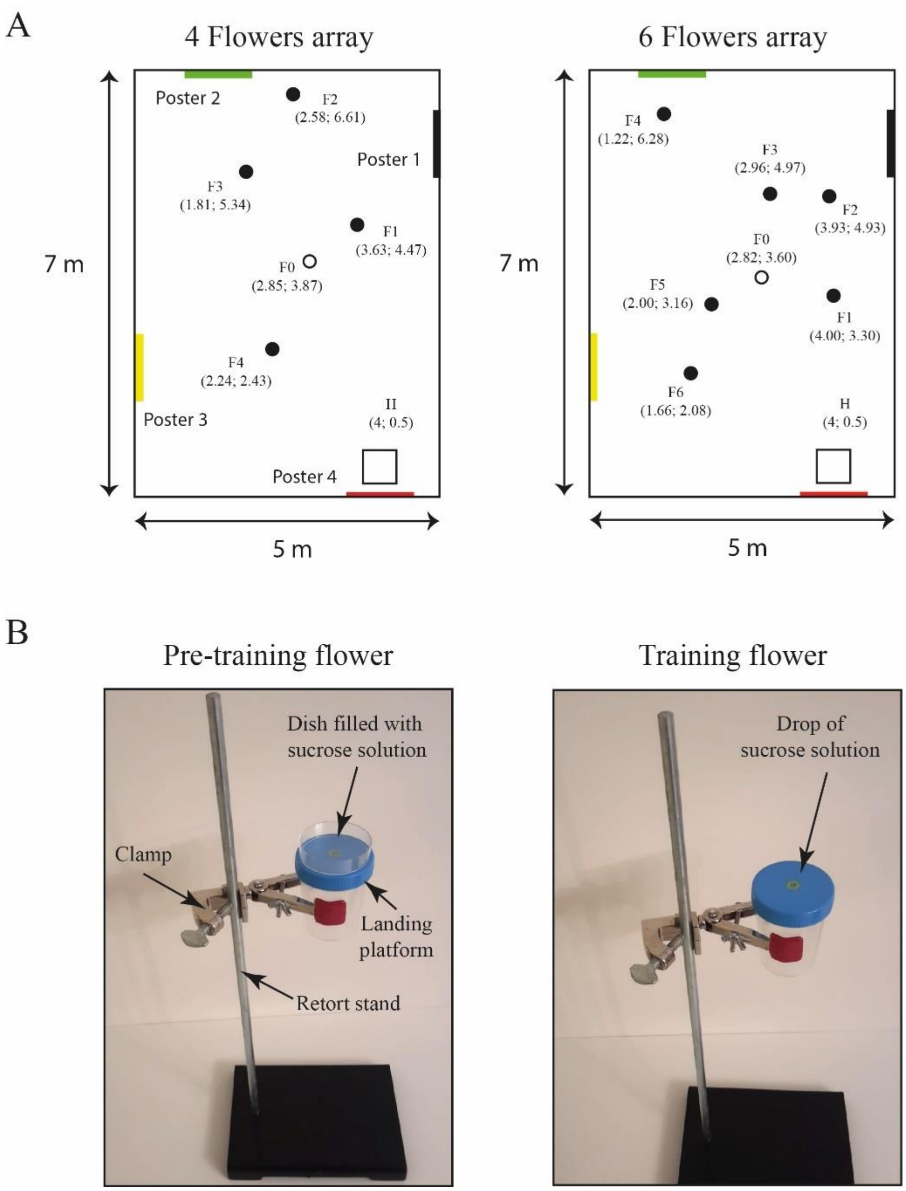


**Figure S1. (A)** Flower arrays. H is the hive, F0 is the pre-training flower, F1-F6 are the experimental flowers and poster 1-4 are the 2D visual landmarks. Number in parentheses are Cartesian coordinates (m). **(B)** Artificial flowers. Flowers consisted of a blue plastic landing platform (diameter = 6cm) sitting on a transparent plastic cylinder (diameter min= 5.5 cm). Each flower was hold 30 cm above ground by a clamp attached to a 50 cm retort stand. A yellow mark in the middle of the landing platform indicated the location of the sucrose reward. Pre-training flowers differed from the experimental flowers by the presence of a petri dish (diameter: 6cm, volume: 110 ml) placed on the landing platform and filled with sucrose to provide bees with *ad libitum* reward. A precise volume of sucrose reward was added manually on the yellow dot of the training flowers using an electronic pipette.

#### Figure S2

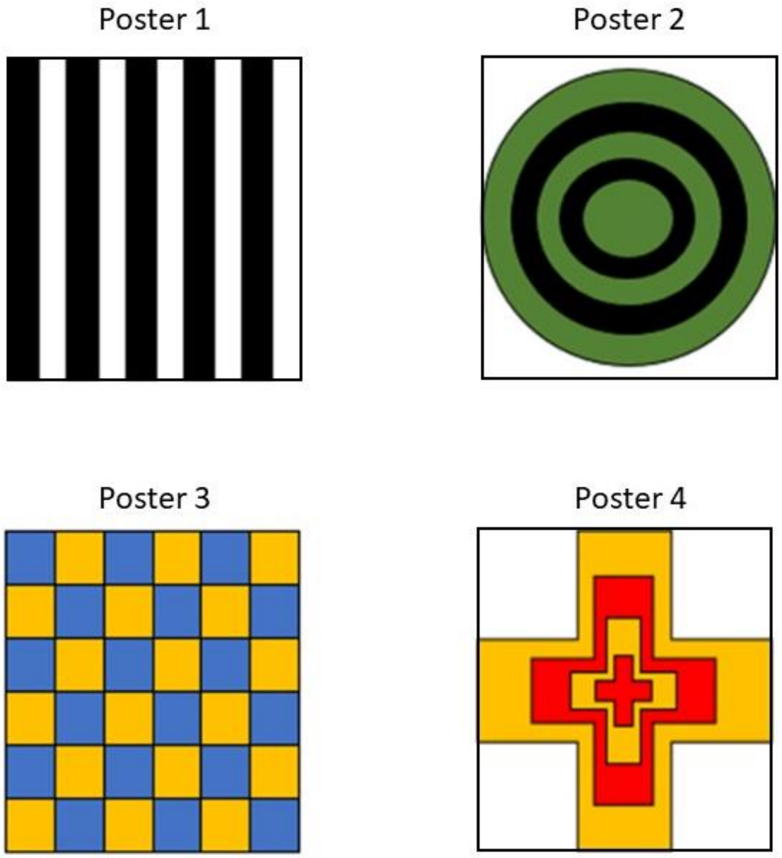


**Figure S2.** Visual appearance of the geometric-patterned posters used in the experiment. Each poster (dimension A0) was positioned on a different wall of the flight room, providing 2D visual landmarks to bees (see precise locations in Fig.S1A)
